# Local protein synthesis of mtIF3 regulates mitochondrial translation for axonal development

**DOI:** 10.1101/2021.01.26.428248

**Authors:** Soyeon Lee, Dongkeun Park, Chunghun Lim, Jae-Ick Kim, Kyung-Tai Min

## Abstract

Mitochondrial initiation factor 3 (mtIF3) binds to and dissociates mitochondrial ribosomes. The mtIF3-small subunit complex then recruits mtIF2, mRNA, and N-formylmethionine-tRNA to initiate mitochondrial translation. Intriguingly, transcripts of the nuclear-encoded mtIF3 gene have been shown present in axonal growth cones; however, the biological function of this compartmentalization remains largely unknown. Here, we demonstrate that brain-derived neurotrophic factor (BDNF) induces local translation of mtIF3 mRNA in axonal growth cones. Subsequently, mtIF3 protein is translocated into axonal mitochondria and promotes mitochondrial translation as assessed by our newly developed bimolecular fluorescence complementation sensor for the assembly of mitochondrial ribosomes. We further show that BDNF-induced axonal growth requires mtIF3-dependent mitochondrial translation in axons. These findings provide new insight into how neurons adaptively control mitochondrial physiology and axonal development via local mtIF3 translation.

## Introduction

Mitochondrial oxidative phosphorylation (OXPHOS) complexes primarily generate ATP essential for cellular function in neuronal cell bodies and neurites. In fact, mitochondria are transported to axons and produce local energy for axon branching, growth cone formation, and axon growth (Rangaraju, Lauterbach, & Schuman, 2019; Spillane, Ketschek, Merianda, Twiss, & Gallo, 2013; Vaarmann et al., 2016). The localized mitochondria also play a significant role in facilitating axonal regeneration after injury (Han, Baig, & Hammarlund, 2016; Lee, Wang, Hwang, Namgung, & Min, 2019; Zhou et al., 2016). Thus, rapid ATP synthesis in response to local energy demand is likely crucial, particularly for polarized neuronal function.

Although active mitochondrial transport to axonal tip has been shown to support local energy needs (Saxton & Hollenbeck, 2012; Sheng, 2017), this may not be sufficient to explain how neurons adaptively regulate mitochondrial function in axons (Niescier, Kwak, Joo, Chang, & Min, 2016). Hence, we reason that additional mechanisms, such as local synthesis of mitochondrial proteins, should contribute to the functional control of axonal mitochondria. Most mitochondrial genes are nuclear-encoded, and once transcribed, their mRNA translation generally occurs in the cell bodies. On the other hand, previous studies have revealed that transcripts of the nuclear-encoded mitochondrial genes can be locally translated in axons (A. Aschrafi et al., 2016; Gale, Aschrafi, Gioio, & Kaplan, 2018; Kaplan, Gioio, Hillefors, & Aschrafi, 2009; Kuzniewska et al., 2020; Shigeoka et al., 2016). Nonetheless, it is still elusive whether any local synthesis of the nuclear-encoded mitochondrial proteins governs mitochondrial function in axons.

In mammalian cells, mitochondria have only two mitochondrial translation initiation factors, mtIF2 and mtIF3 (Smits, Smeitink, & van den Heuvel, 2010). Interestingly, translatome analyses have revealed mtIF3 translation in axon growth cone (Shigeoka et al., 2016), suggesting a possible role of local mtIF3 synthesis in regulating axonal mitochondrial translation. mtIF3 regulates the dynamics of ribosome association on mitochondrial mRNAs. mtIF3 catalyzes the dissociation of mitochondrial ribosomes (mitoribosomes) into large and small subunits while blocking any premature binding of the large subunit (Christian & Spremulli, 2009; Koc & Spremulli, 2002). mtIF2 and N-formylmethionine-tRNA bind weakly to the small subunit in the absence of mRNA, but mtIF3 facilitates mRNA binding to the small subunit so that a start codon can be correctly positioned at P-site (Christian & Spremulli, 2009; Smits et al., 2010).

Given the critical role of mtIF3 in mitochondrial translation initiation (Rudler et al., 2019), it is plausible that locally synthesized mtIF3 may regulate mitochondrial translation in developing axons to support ATP synthesis and relevant physiology. Studies on mitochondrial translation in live cells, however, have been hampered by a lack of appropriate tools. Here, we have developed a molecular sensor that visualizes mitochondrial translation activity using the bimolecular fluorescence complementation (BiFC) between a specific pair of mitoribosome proteins. In conjunction with additional transgenic reporters for functional imaging, this new tool has led us to test the hypothesis above and validate the significance of local mtIF3 translation in mitochondrial physiology and axonal growth.

## Results

### BDNF induces local protein synthesis of mtIF3 in axon growth cone

We first confirmed that mtIF3 mRNAs were present in both cell bodies and axons of primary hippocampal neurons (Figure 1A), consistent with a previous report (Shigeoka et al., 2016). To examine whether locally translated mtIF3 proteins translocate into mitochondria, we generated a transgene that expresses fluorescent mtIF3 proteins fused to the photo-convertible Dendra2 along with N-terminal palmitoylation sequence (mtIF3-Dendra2) (Lee et al., 2019; Wang et al., 2016) and mtIF3 untranslated regions (UTRs) (5’UTR_mtIF3_-mtIF3-Dendra2-3’UTR_mtIF3_). As expected, the coding sequence (CDS) of mtIF3 led to mitochondrial localization of mtIF3-Dendra2 fusion likely due to its mitochondrial targeting sequence (Figure 1-figure supplement 1). The fluorescent mtIF3-Dendra2 proteins in axons were irreversibly photo-switched from green to red using 405 nm illumination, and then newly translated mtIF3-Dendra2 proteins with green fluorescence were measured by analyzing time-lapse images taken every 5 minutes for 90 minutes. Several studies have suggested that many nuclear-encoded mitochondrial proteins might be synthesized in response to local energy demand (Gale et al., 2018; Kaplan et al., 2009; Kuzniewska et al., 2020; Shigeoka et al., 2016). This prompted us to examine whether BDNF treatment in axonal growth cone enhances the local translation of the mtIF3-Dendra2 fusion reporter. Indeed, kymograph analyses revealed that BDNF treatment elevated the newly synthesized mtIF3-Dendra2 signals in axonal mitochondria (Figure 1B-C). A translation inhibitor, anisomycin, blocked the de novo synthesis of reporter proteins, validating that BDNF treatment triggers the local synthesis of mtIF3 proteins in axons and promotes their translocation into axonal mitochondria.

**Figure 1.**
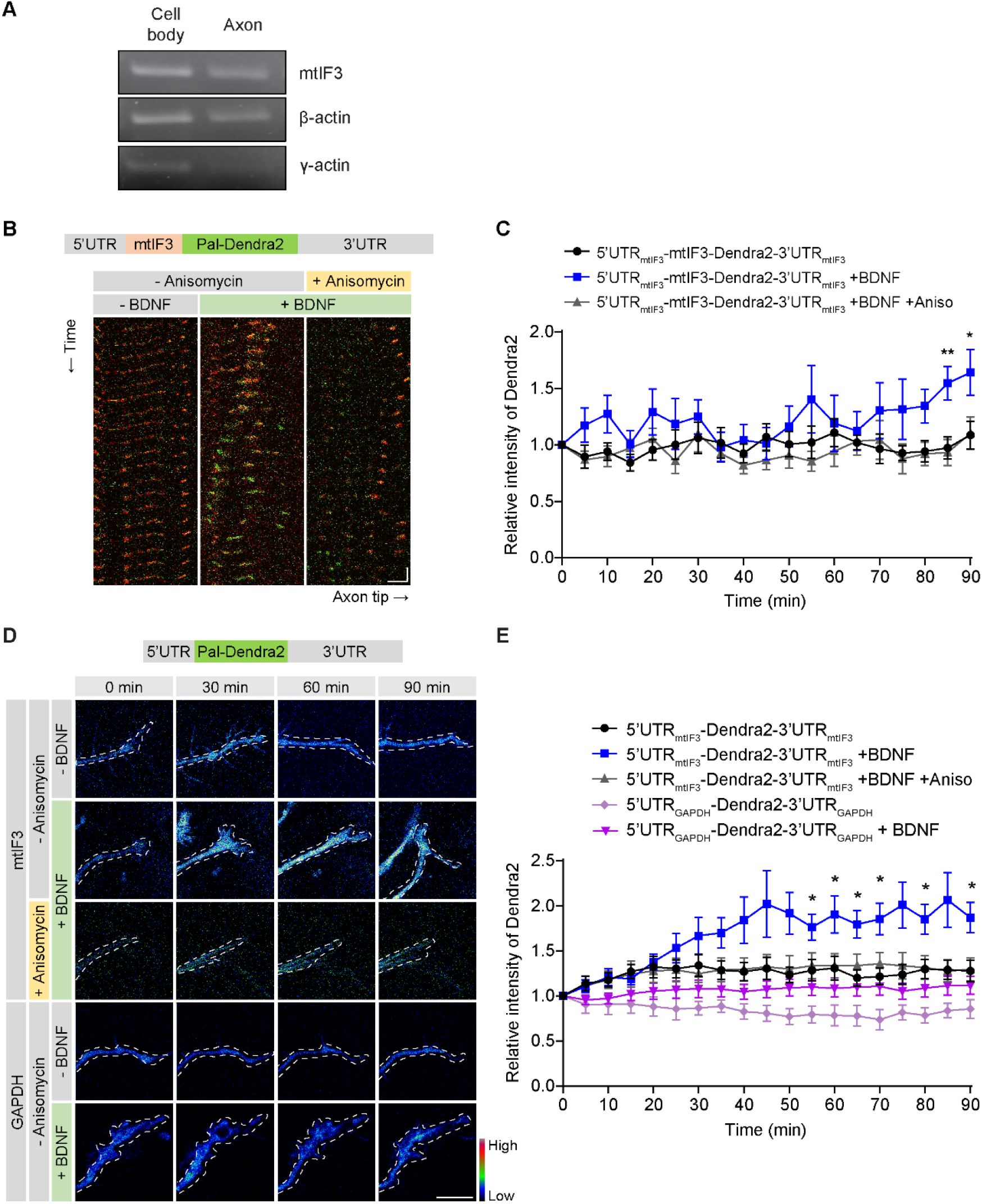
BDNF induces local protein synthesis of mtIF3 in the axon growth cone. (**A**) mtIF3 mRNAs were detected in both cell bodies and axons of primary hippocampal neurons at DIV4. RNA samples were purified from the isolated lysates of cell bodies and axons. RT-PCR was performed using each pair of gene-specific primers. (**B**) Kymographs of newly synthesized mtIF3-Dendra2 fusion proteins in axons. Primary hippocampal neurons were transfected with an expression vector for the photoconvertible mtIF3-Dendra2 protein at DIV3, and the fluorescent intensity of mtIF3-Dendra2 fusion was analyzed at DIV4 (horizontal scale bar, 5 μm; vertical scale bar, 5 minutes). The existing mtIF3-Dendra2 was first photoconverted from green to red over the 20 μm path from the axon tip. Images were then taken at 5-minute intervals for 90 minutes. Where indicated, BDNF (30 ng/ml) and anisomycin (20 μM) were added at a 0-minute timepoint. (**C**) Quantification of newly synthesized mtIF3-Dendra2 proteins. The green fluorescence was measured from mitochondria at the very end of the axonal tip. The relative intensity at each time point was calculated by normalizing to that at a 0-minute timepoint. Data represent mean ± SEM (N = 6-8 axons from 3 independent experiments). \**P* < 0.05, \*\**P* < 0.01, as determined by two-way repeated-measures ANOVA with Dunnett’s multiple comparisons test. (**D**) Pseudo-color images of locally synthesized Dendra2 reporter in the axonal tip. Primary hippocampal neurons were transfected with an expression vector for the photoconvertible Dendra2 reporter harboring the indicated UTRs at DIV3. The newly synthesized Dendra2 reporter was analyzed at DIV4 (scale bar, 10 μm) similarly as in panel B. (**E**) Quantification of newly synthesized Dendra2 reporters in the axonal tip after drug treatment. Data represent mean ± SEM (N = 6-10 axons from 3 independent experiments). \**P* < 0.05, as determined by two-way repeated-measures ANOVA with Dunnett’s multiple comparisons test.

Interestingly, mtIF3 3’UTR contains a consensus motif (CTCCCATC) shared by axon-enriched mRNAs (Shigeoka et al., 2016). We thus generated two additional translation reporters encoding the fluorescent Dendra2 along with UTRs from mtIF3 (5’UTR_mtIF3_-Dendra2-3’UTR_mtIF3_) or GAPDH (5’UTR_GAPDH_-Dendra2-3’UTR_GAPDH_). BDNF treatment gradually increased the fluorescence of newly synthesized Dendra2 from the mtIF3 UTR reporter at the axonal tip, whereas anisomycin treatment suppressed it (Figure 1D-E). We detected no significant changes in the fluorescence from the control GAPDH UTR reporter upon BDNF treatment. Together, these results indicate that mtIF3 proteins are locally synthesized in axon growth cones and translocate into mitochondria in response to BDNF. In addition, mtIF3 UTRs likely support the axonal transport of mtIF3 mRNAs and their BDNF-induced translation in axons, regardless of mitochondrial targeting of the translation products.

### Mito-riboBiFC detects translation-dependent assembly of mitoribosomes

We hypothesized that BDNF-induced local translation of mtIF3 proteins might be involved in regulating mitochondrial translation in developing axon tip. To overcome possible limitations in the biochemical assessment of mitochondrial translation in axons, we devised a new strategy to visualize mitochondrial translation in live cells using BiFC (Hu, Chinenov, & Kerppola, 2002) which was previously used for the visualization of cytoplasmic ribosomal subunit joining (Al-Jubran et al., 2013). This was based on the physical proximity of mitochondrial ribosomal protein L2 (MRPL2) and mitochondrial ribosomal protein S6 (MRPS6) at the inter-subunit bridge of 55S mitoribosome (Amunts, Brown, Toots, Scheres, & Ramakrishnan, 2015) (Figure 2A). We took advantage of this adjacent localization of the two MRPs as a BiFC pair to visualize mitoribosome assembly during translation. In detail, we split a fluorescent protein mVenus into N-terminal (VN, 1-172 amino acids) and C-terminal fragments (VC, 155-238 amino acids). Then we fused these mVenus fragments to the C-termini of MRPS6 (S6-VN) and MRPL2 (L2-VC), respectively. The co-expression of S6-VN and L2-VC in Neuro2A cells generated mVenus fluorescent signals exclusively in mitochondria (Figure 2B). On the other hand, another pair of MRPs positioned distantly from each other in mitoribosomes showed relatively weak fluorescent signals (Figure 2A-D, MRPS16 and MRPL50) even though the latter BiFC pair were expressed comparably to S6-VN and L2-VC (Figure 2B-D).

Unlike cytoplasmic ribosomes, two mitoribosome subunits could be assembled in the absence of translating mRNAs (Smits et al., 2010). We thus asked if the BiFC signals from the S6-VN and L2-VC pair would depend on the translation of mitochondrial mRNAs. When we treated puromycin to dissociate translating ribosome subunits from mRNAs, the BiFC signals were reduced only by 8% compared to the vehicle control (Figure 2E and F), indicating the mRNA-independent assembly of the BiFC pair. The average cytoplasmic translation rate is six amino acids per second (Ingolia, Lareau, & Weissman, 2011), and the longest transcript mt-ND5 mRNA is 1824 bp. In contrast, the fluorescence signals from the BiFC pair become detectable 10 minutes after complementation (Robida & Kerppola, 2009). Considering that the translation of individual mitochondrial mRNAs could be completed in less than 2 minutes, it is likely that the translating mitoribosome dissociates from mRNAs before the chromophore maturation. We thus reasoned that the stabilization of translating mitoribosome would better visualize their mRNA-dependent emission of the complemented fluorescence signals. Indeed, chloramphenicol (CA), an inhibitor of the peptide bond formation at mitoribosome E-site, markedly increased the BiFC signals 60 minutes after treatment, whereas puromycin pre-treatment blocked the CA effects (Figure 2E-F). Accordingly, we concluded that CA-induced BiFC signals would represent actively translating mitoribosome and designated our new tool for visualizing mitochondrial translation as mito-riboBiFC (Figure 2-figure supplement 1).

**Figure 2.**
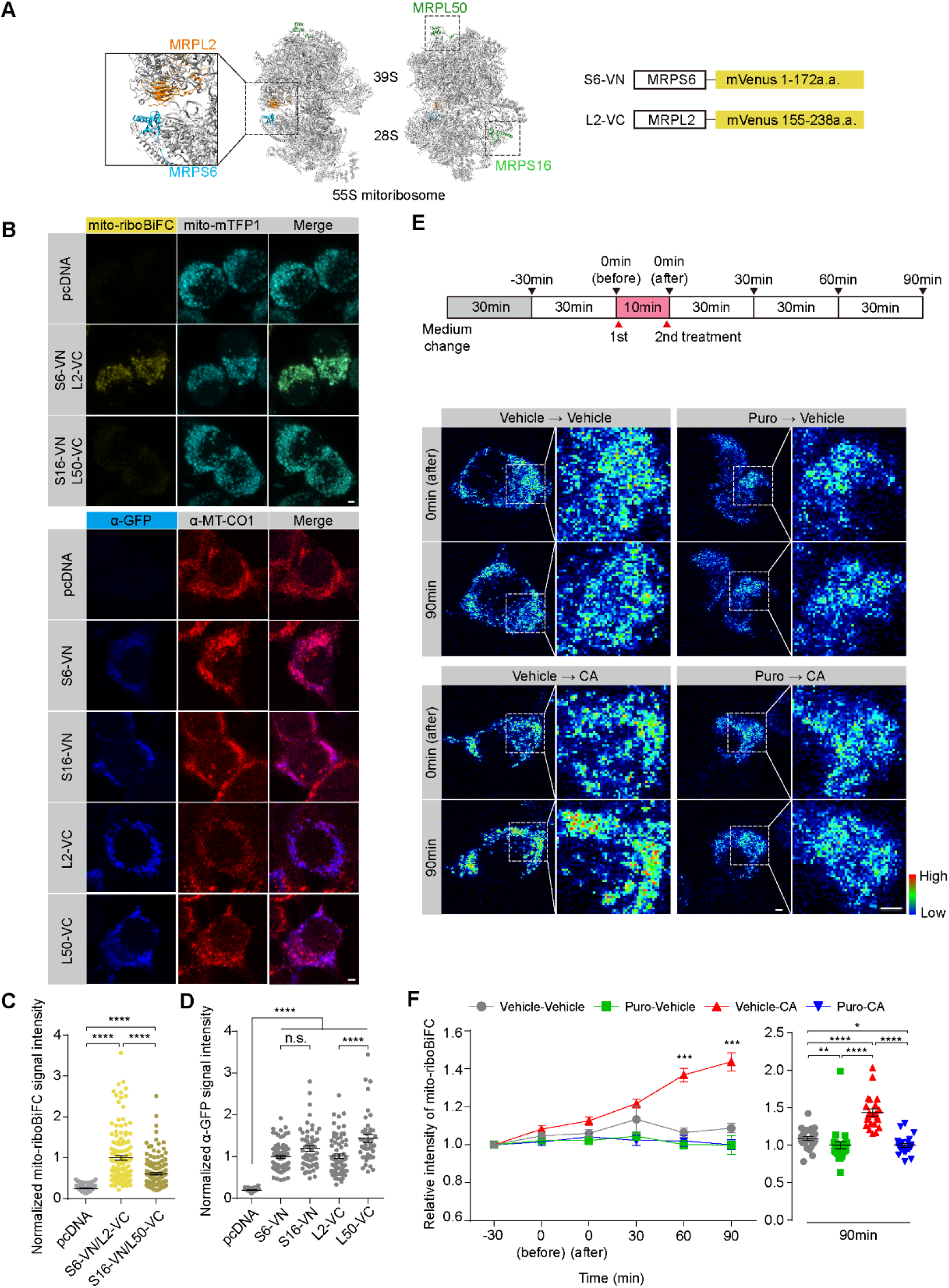
Mito-riboBiFC detects translation-dependent assembly of mitoribosomes. (**A**) Schematic design of mito-riboBiFC. Mitochondrial ribosomal proteins MRPL2 and MRPS6 were used as a BiFC pair for the mito-riboBiFC and illustrated with porcine 55S mitoribosome cryo-EM structure (Greber et al., 2015). MRPS16 and MRPL50 served as a negative control. (**B**) Representative images of mito-riboBiFC (top) and anti-GFP antibody staining (bottom) in Neuro2A cells. Mito-mTFP1 and MT-CO1 were used as mitochondrial markers (scale bar, 2 μm). (**C**) Quantification of the fluorescent mito-riboBiFC signals in panel B. Data represent mean ± SEM (N = 100-143 cells from 3 independent experiments). \*\*\*\**P* < 0.0001, as determined by aligned ranks transformation ANOVA with Wilcoxon rank-sum test. (**D**) Quantification of At-GFP signal intensity in panel B. Data represent mean ± SEM (N = 41-76 cells from 3 independent experiments). n.s., not significant; \*\*\*\**P* < 0.0001, as determined by aligned ranks transformation ANOVA with Wilcoxon rank-sum test. (**E**) Pseudo-color images of mito-riboBiFC after sequential treatment of puromycin (Puro) and chloramphenicol (CA) (scale bar, 2 μm). (**F**) Quantification of the mito-riboBiFC signals in panel E. Line plot shows the intensity changes of mito-riboBiFC. A Vehicle-CA group was compared with other groups at each time point for the statistical test (left panel). Dot plot displays the relative intensity of the mito-riboBiFC 90 minutes after CA treatment (1^st^ vehicle, water; 2^nd^ vehicle, ethanol). Data represent mean ± SEM (N = 22-26 cells from 3 independent experiments). Aligned ranks transformation ANOVA detected significant interaction effects of Puro and CA on the mito-riboBiFC intensity at the 90-minute time point (*P* < 0.0001). \*\**P* < 0.01, \*\*\*\**P* < 0.0001, as determined by Wilcoxon signed-rank test (left panel) or Wilcoxon rank-sum test (right panel).

### Locally synthesized mtIF3 promotes mitochondrial translation in axon growth cone

To test whether locally synthesized mtIF3 facilitates mitochondrial translation in the axon growth cone, we manipulated mtIF3 expression by transient transfections and examined their effects on the mito-riboBiFC signals in axons. We first confirmed mtIF3 depletion using short hairpin RNA (shRNA) or transgenic mtIF3 overexpression in NIH/3T3 cells (Figure 3-figure supplement 1). Next, we cultured hippocampal neurons on a microfluidic device to separate axons from cell bodies (Taylor et al., 2005) and treated all the drugs only in the axonal channel to induce or block local translation (Figure 3A-B). The degree of mitochondrial translation was subsequently quantified using CA-induced changes in the intensity of mito-riboBiFC (Figure 2-figure supplement 1). Notably, neurons expressing control shRNA exhibited BDNF-induced BiFC signals in the axon growth cone. However, the treatment of a translation inhibitor cycloheximide (CHX) completely blocked the increment of BiFC signals upon BDNF treatment, indicating that BDNF-induced local protein synthesis promotes mitochondrial translation. Importantly, mtIF3 depletion abolished the BDNF-induced mito-riboBiFC signals (Figure 3C-D), suggesting that locally synthesized mtIF3 is necessary for facilitating mitochondrial translation upon BDNF treatment.

**Figure 3.**
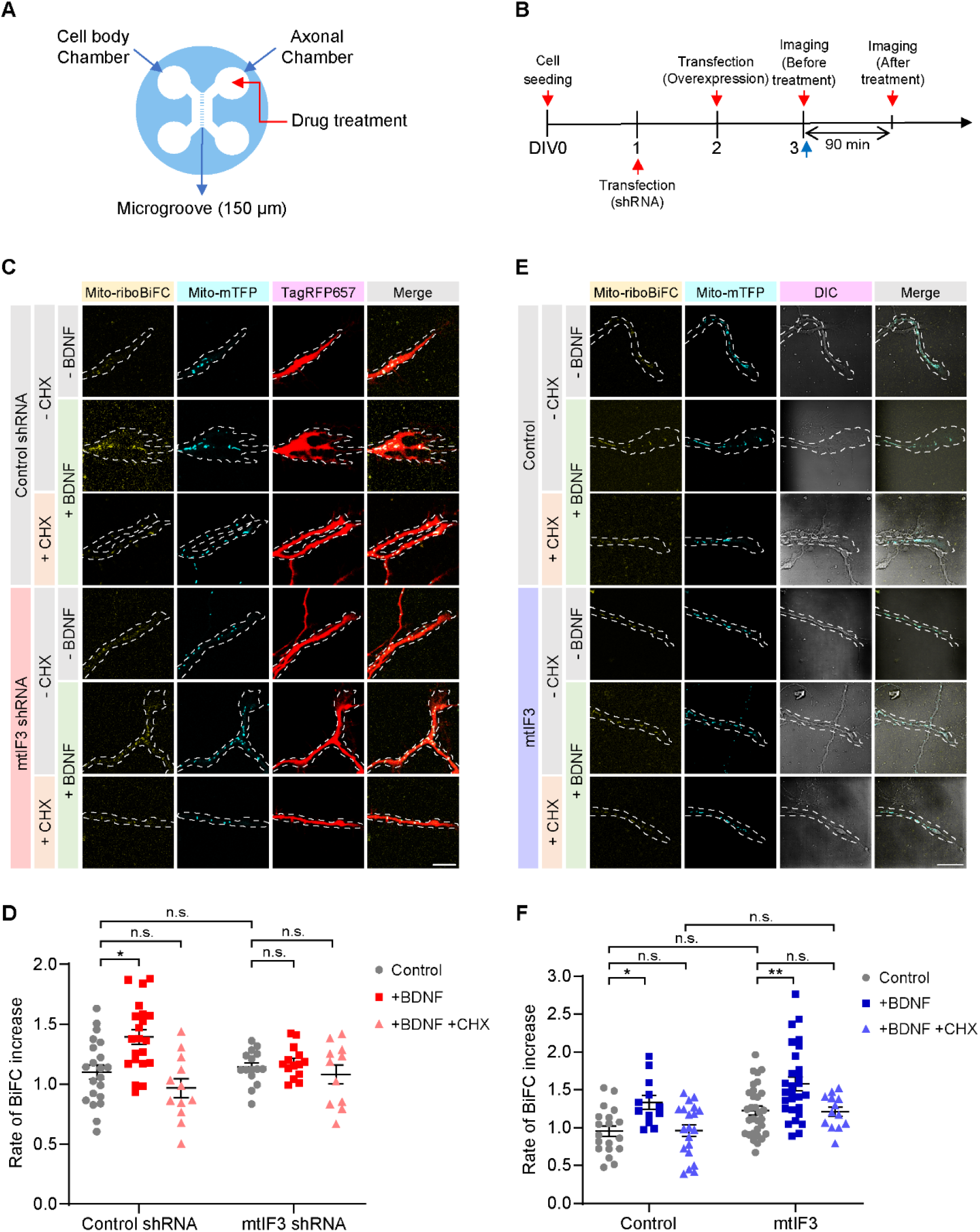
Local protein synthesis of mtIF3 is necessary for mitochondrial translation in the axonal growth cone. (**A**) Schematic illustration of a microfluidic device. Primary hippocampal neurons were seeded into the cell body chamber. Growing axons reached the other side of the device through the microgroove. To locally stimulate axons, drugs were treated to the axonal chamber. (**B**) Timeline for mito-riboBiFC experiments. After cell seeding, shRNA and overexpression vectors were transfected at DIV2 and DIV3, respectively. Images were sequentially taken before and after drug treatment. (**C, D**) Visualization of mitochondrial translation in mtIF3-depleted axon growth cones by mito-riboBiFC. Mitochondria were marked by mitochondria-targeted mTFP 1. Transfection of shRNA was confirmed by TagRFP657 expression (scale bar, 20 μm). Mito-riboBiFC was quantified, and the rate of BiFC increase upon drug treatment was measured. Five mitochondria per axon were analyzed. Data represent mean ± SEM (N = 11-21 axons from 3 independent experiments). Aligned ranks transformation ANOVA detected significant interaction effects of mtIIF3 depletion and BDNF on the BiFC rate (*P* = 0.0180). n.s., not significant; \**P* < 0.05, as determined by Wilcoxon rank-sum test (BDNF) or two-way ANOVA with Tukey’s multiple comparisons test (BDNF+CHX). (**E, F**) Representative images of mito-riboBiFC in mtIF3-overexpressing axon growth cones (scale bar, 20 μm). Mito-riboBiFC signals were analyzed before and after the CA treatment. Five mitochondria per axon were analyzed. Data represent mean ± SEM (N = 12-31 axons from 4 independent experiments). Two-way ANOVA detected no significant interaction effects of mtIF3 overexpression and drug treatments. n.s., not significant; \**P* < 0.05, \*\**P* < 0.01, as determined by Tukey’s multiple comparisons test.

To assess whether mtIF3 overexpression elevates the mitochondrial translation in the absence of BDNF, we transfected primary hippocampal neurons with a mtIF3 overexpression vector at DIV2 and measured any change in the mito-riboBiFC signals. Neither the baseline nor BDNF-induced mito-riboBiFC signals were significantly affected by mtIF3 overexpression under our experimental conditions. Considering that our transgene for mtIF3 overexpression included mtIF3 UTRs, we reason that local synthesis of mtIF3 would be tightly regulated at post-transcript levels, and the axonal abundance of mtIF3 mRNAs may not be limiting for BDNF-induced mitochondrial translation. Together, these results support that BDNF-induced local protein synthesis of mtIF3 leads to enhanced mitochondrial translation in the axon growth cone.

### mtIF3-dependent mitochondrial translation elevates ATP generation in growing axons

Next, we questioned whether locally translated mtIF3 would control mitochondrial physiology in developing axons. To this end, we employed mito-ATeam1.03, a genetically encoded FRET sensor for mitochondrial ATP (Imamura et al., 2009). CA treatment to primary hippocampal neurons expressing mito-ATeam1.03 reduced the intensity of the FRET signals, indicating that mitochondrial ATP generation requires mitochondrial translation (Figure 4-figure supplement 1). We further found that BDNF treatment elevated mitochondrial ATP levels in the axon growth cone, whereas blocking local translation by CHX nullified the BDNF effects (Figure 4A-B). mtIF3 depletion also blunted BDNF-induced increase in mitochondrial ATP levels, yet it negligibly affected the baseline ATP levels (Figure 4A-B). We observed no significant effects of mtIF3 overexpression on mitochondrial ATP levels in axons regardless of BDNF treatment (Figure 4C-D), consistent with mtIF3 effects on mitochondrial translation in axons. These results support our model that BDNF-induced local synthesis of mtIF3 promotes mitochondrial translation and elevates ATP generation in axonal mitochondria, thereby fulfilling local energy demand in the developing axons.

**Figure 4.**
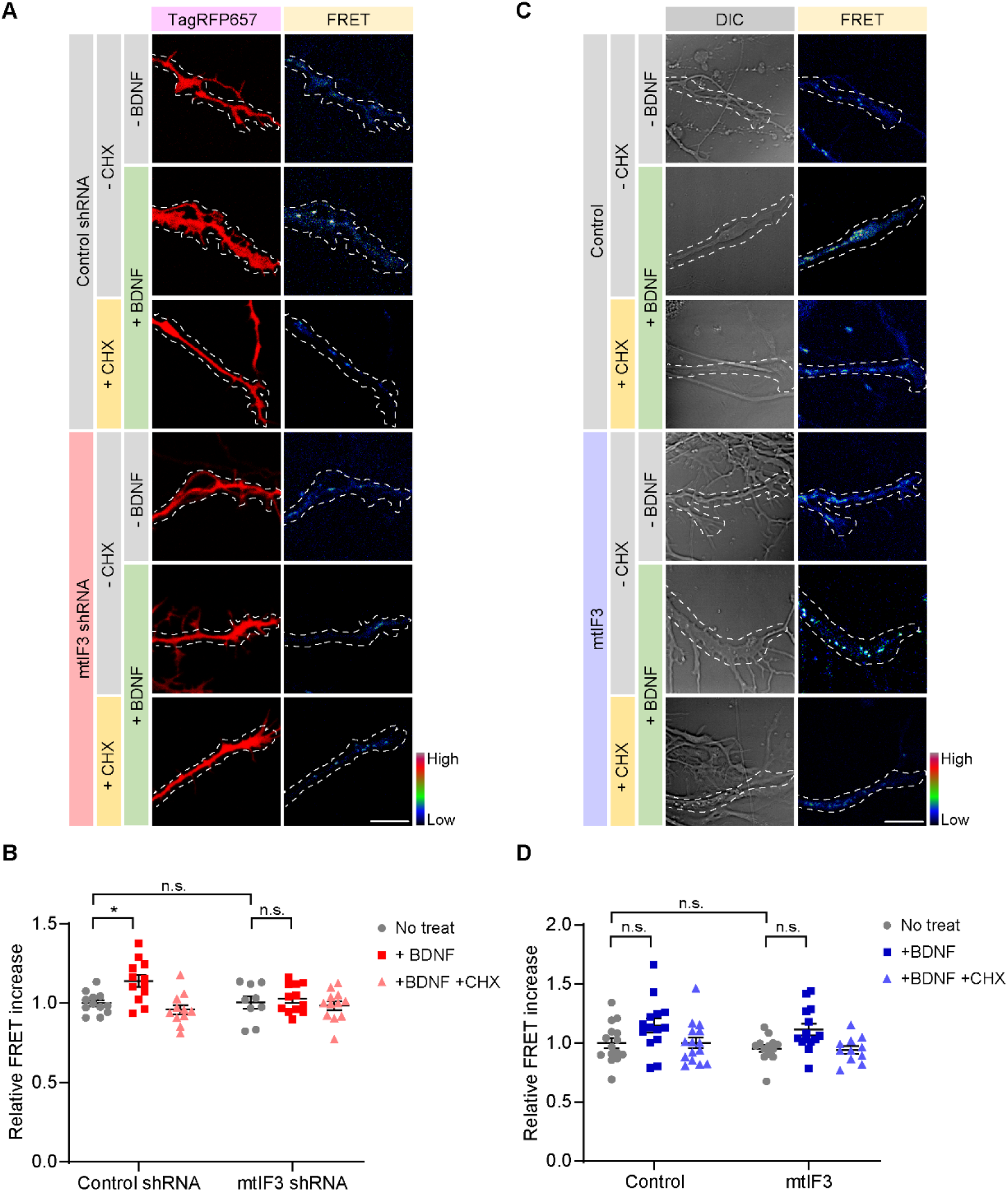
mtIF3-dependent mitochondrial translation elevates ATP generation in the axonal growth cone. (**A, C**) ATP levels in axonal mitochondria were measured using genetically encoded ATP indicator mito-ATeam1.03. FRET signals were shown in the pseudo-color image. Expression of shRNA was confirmed by TagRFP657 expression (scale bar, 10 μm). (**B, D**) Quantification of the relative FRET intensity in mtIF3-depleted or mtIF3-overexpressing axons. FRET signals were measured by comparing the ratio before and after chemical treatments. Five mitochondria per axon were analyzed. Data represent mean ± SEM (N = 9-15 axons from 4-5 independent experiments). Two-way ANOVA detected significant effects of BDNF, but not of BDNF+CHX, on the FRET signals (*P* = 0.0124 in panel B; *P* = 0.0020 in panel D). n.s., not significant; \**P* < 0.05, as determined by Tukey’s multiple comparisons test.

### Axonal development requires mtIF3-dependent mitochondrial translation in growing axons

To determine whether local mitochondrial translation indeed impacts axonal growth, we applied CA to either cell bodies or axons of primary hippocampal neurons cultured in a microfluidic device (Figure 5A). BDNF was subsequently added to the axonal chamber, and BDNF-induced axon growth was quantified accordingly. We found that selective inhibition of mitochondrial translation in axons, but not in cell bodies, suppressed BDNF-induced axon extensions (Figure 5B-C). These data demonstrate that rapid axon extension by this trophic factor requires local mitochondrial translation in axons. Given that locally synthesized mtIF3 regulates the mitochondrial translation in axons, we reasoned that mtIF3 depletion would impair axonal extension. Indeed, transient overexpression of mtIF3 shRNA remarkably shortened axonal length compared to control shRNA (Figure 5D-E). Moreover, mtIF3 depletion silenced BDNF effects on axon development. We observed that mtIF3 overexpression negligibly affected axon growth regardless of BDNF treatment (Figure 5-figure supplement 1), consistent with its lack of any significant effects on mitochondrial translation and ATP generation in axons. Together, our findings validate that mtIF3-dependent mitochondrial translation in axons plays a critical role in axonal development.

**Figure 5.**
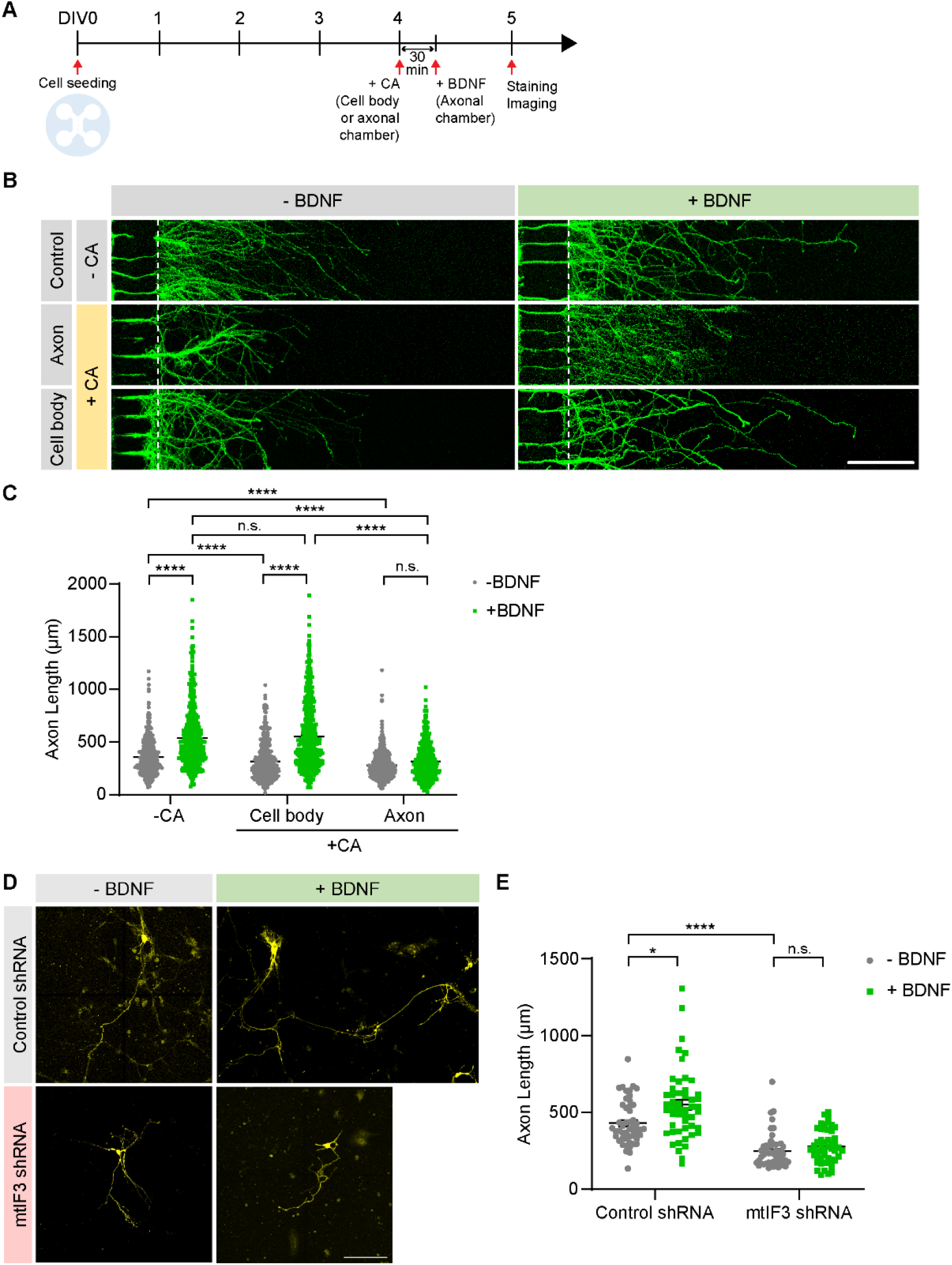
Axonal development requires mtIF3-dependent mitochondrial translation in growing axons. (**A**) Timeline for the experiment. Primary hippocampal neurons were cultured on a microfluidic device. At DIV4, chloramphenicol (CA) was added to either the cell body or axonal chamber. After 30 minutes, BDNF was added to the axonal chamber. Neurons were stained and imaged at DIV5. (**B**) Representative images of axons. Axons were marked by Tau-1 immunostaining (scale bar, 200 μm). (**C**) The axonal length was measured from the exit border of microgrooves (dotted lines). Data represent mean ± SEM (N = 422-745 axons from 4-5 independent experiments). n.s., not significant; \*\*\*\**P* < 0.0001, as determined by Aligned ranks transformation ANOVA with Wilcoxon rank-sum test. (**D**) Representative images of mtIF3-depleted or BDNF-treated primary hippocampal neurons. mtIF3 depletion impaired the extension of the axon growth cone. Hippocampal neurons were first transfected with mtIF3 shRNA and then treated with BDNF at DIV3. The axonal length was measured at DIV5 (scale bar, 100 μm). (**E**) quantification of axon length in panel D. Data represent mean ± SEM (N = 50 neurons from 5 independent experiments). Aligned ranks transformation ANOVA detected significant interaction effects of mtIIF3 depletion and BDNF on the axon length (*P* = 0.0375). n.s., not significant; \**P* < 0.05, \*\*\*\**P* < 0.0001, as determined by Wilcoxon rank-sum test

## Discussion

Local protein synthesis is a distinctive feature in neurons that have highly polarized neurites, which is indispensable for the maintenance of axonal or dendritic structures and functions such as neurite development, the guidance of growth cone, synaptic transmission, synaptic plasticity, branch formation, and regeneration (Hafner, Donlin-Asp, Leitch, Herzog, & Schuman, 2019; Jung, Yoon, & Holt, 2012; Shigeoka et al., 2019). The gene ontology analyses revealed high enrichment of synaptic proteins, cytoskeletal proteins, and ribosomal proteins in axonal translatome (Gumy et al., 2011; Shigeoka et al., 2016). Interestingly, it has been also identified that transcripts of nuclear-encoded mitochondrial proteins are abundant in developing and mature axons (A. Aschrafi et al., 2016; Gale et al., 2018; Kaplan et al., 2009; Kuzniewska et al., 2020; Shigeoka et al., 2016), implicating their local translation in sustaining mitochondrial function and axonal viability. Nonetheless, only a few studies have documented that local translation of nuclear-encoded mitochondrial proteins can affect mitochondrial function and axonal survival (Armaz Aschrafi, Natera-Naranjo, Gioio, & Kaplan, 2010; Hillefors, Gioio, Mameza, & Kaplan, 2007; Natera-Naranjo et al., 2012; Yoon et al., 2012).

Here, we demonstrate that nuclear-encoded mtIF3 is locally translated in developing axons, thereby promoting axonal mitochondrial translation as assessed by our newly developed mito-riboBiFC sensor. Many studies have demonstrated that stationary mitochondria in axons fuel spatially restricted boundaries (Rangaraju et al., 2019; Spillane et al., 2013), but what remains unsolved is how these stationary mitochondria are supported and maintained in the long-term. Our results suggest that mitochondrial proteins may be replenished by enhanced mitochondrial translation via local protein synthesis in axons. We observed that the mtIF3 depletion cancels out the upregulation of mitochondrial translation and ATP production upon BDNF stimulation. Lack of this local translation and adaptive control of mitochondrial function limits axonal development, validating its critical role in neuronal physiology. However, we also observed that the overexpression of mtIF3 *per se* did not affect mitochondrial functions. It has been recently shown that mitochondrial translation is synchronized and unidirectionally controlled by cytosolic translation (Couvillion, Soto, Shipkovenska, & Churchman, 2016). Our observation consistently implicates that enhanced mitochondrial functions for local energy demanding are accomplished by not the mitochondrial translation alone but the simultaneous cytosolic translation. Therefore, our findings suggest that local translation in axons can be a crucial mechanism by which mitochondrial translation is regulated in mammalian neurons.

In the past decade, much effort has been made to develop the tools for measuring or observing mitochondrial translation. For instance, biochemical detection of newly synthesized mitochondrial proteins has been widely used for studying mitochondrial translation (Barsh et al., 2015; Chatenay-Lapointe & Shadel, 2011; Park, Lee, & Min, 2020; Richter-Dennerlein et al., 2016; Richter, Lahtinen, Marttinen, Suomi, & Battersby, 2015). However, a lack of appropriate imaging tools for mitochondrial translation has hindered assessing this subcellular event at single-cell levels. A recent study visualized mitochondrial translation using a non-canonical amino acid labeling *in situ* (Estell, Stamatidou, El-Messeiry, & Hamilton, 2017). This method allows the detection of mitochondrial translation at a singlecell resolution, but its application is limited to fixed cells. Our study developed a new method designated as mito-riboBiFC to monitor mitochondrial translation in live cells. Mito-riboBiFC enables us to investigate mitochondrial translation on distinct spatiotemporal scales. Accordingly, it will be of great interest to determine how mitochondrial translation is regulated depending on their subcellular location or mitochondrial dynamics, especially in neurons where subcellular environment and energetic needs are spatially distinct.

Nonetheless, mito-riboBiFC has some limitations that should be improved in the future. These include relatively slow maturation kinetics of the mito-riboBiFC. Mitochondria are highly dynamic and heterogeneous in terms of their transport, membrane potential, and biogenesis. These mitochondrial events can occur on a relatively short timescale (e.g., a few seconds or minutes), compared to the folding and maturation time of the BiFC complex (Rose, Briddon, & Holliday, 2010). The employment of a new chromophore in BiFC imaging should improve the current temporal resolution of our mito-riboBiFC, better visualizing the rapid change in mitochondrial translation according to diverse mitochondrial dynamics.

In conclusion, our results provide new insights into understanding the adaptive regulation of mitochondrial physiology via local protein synthesis of a nuclear-encoded mitochondrial translation factor during axonal development. New imaging tools for the mitochondrial function should further dissect the molecular mechanisms underlying the spatiotemporal control of mitochondria physiology and hint at novel therapeutic strategies to treat relevant neurodevelopmental diseases.

## Materials and methods

### Animals

Pregnant mice (C57BL/6J, Hyochang Science, Korea) were used for primary hippocampal neuron culture. All experimental procedures were conducted in accordance with protocols approved by Institutional Animal Care and Use Committee of Ulsan National Institute of Science and Technology (UNIST).

### Cell culture

#### Primary hippocampal neurons

Primary hippocampal neuron culture was processed as follows. In brief, hippocampi were dissected from E18 mouse embryos and they were washed with HBSS (Invitrogen). Hippocampi were digested by 0.025% trypsin (Invitrogen) and washed with trituration media (90% of Dulbecco Modified Eagle Medium and 10% fetal bovine serum, Invitrogen). Dissociated cells were seeded onto 50 μg/ml of poly-D-lysine (Sigma) coated culture dishes or coverslips. After settlement of cells, neurons were maintained with neuronal culture media, which consists of Neurobasal media, GlutaMax, B27, and penicillin-streptomycin (Invitrogen). Neurons were transfected with lipofectamine 2000 (Invitrogen).

#### Cell lines

Neuro2A cell line was used for mito-riboBiFC experiments and purchased from ATCC. Neuro2A cells were maintained in culture media, which consists of Dulbecco Modified Eagle Medium and 10% fetal bovine serum, and 1% penicillin-streptomycin (Invitrogen). Using mycoplasma detection kit (Takara, 6601), we confirmed no contamination in Neuro2A cell line. PEI (Polysciences, 23966-1) or lipofectamine 2000 (Invitrogen) were used for transfecting constructs into Neuro2A cells. NIH/3T3 was purchased from ATCC and used for the modulation of mtIF3 expression. Cells were maintained in culture media, which consists of Dulbecco Modified Eagle Medium and 10% calf serum, and 1% penicillin-streptomycin (Invitrogen). Metafectene (Biontex) was used for the transfection of mtIF3 constructs.

#### Separation of cell bodies and axons

To isolate lysate of cell bodies and axons separately, neurons were seeded on the 6-well inserts with 3 μm pore size (SPL Life Sciences). Samples of cell bodies and axons were collected by scrapping the upper and bottom side of inserts. To treat cell bodies or axons separately with drugs, neurons were placed on microfluidic devices (Xona Microfluidics). Microfluidic devices were attached to glass bottom dish (In Vitro Scientific, D60-30-1.5) for live cell imaging or 22 mm square coverslips (Globe Scientific, 1404-15) for fixed samples. 30 ng/ml of BDNF (Sigma), 50 μg/ml of chloramphenicol (Sigma), 20 μM of anisomycin (Sigma), and 100 μg/ml of cycloheximide (Sigma) were used for drugs treatment.

#### Vector preparation

For local protein synthesis assay, *pDendra2-C* vector (Evrogen) was modified: *5’UTR of mtIF3-2xPal-Dendra2-3’UTR of mtIF3, 5’UTR of mtIF3-CDS of mtIF3-2xPal-Dendra2-3’UTR of mtIF3, and 5’UTR of GAPDH-2xPal-Dendra2-3’UTR of GAPDH*. To block the effect of diffusion, two repeats of palmitoylation sequence was added. For mito-riboBiFC assay, *pcDNA6/V5-HisA* (Invitrogen) plasmid was modified: Neuro2A cDNA sequence of Mouse *Mrps6* (NM_080456.1), *Mrpl2* (NM_025302.4), *Mrps16* (NM_025440.3), and *Mrpl50* (NM_178603.4) were used to generate *MRPS6-VN172, MRPL2-VC155, MRPS16-VN172*, and *MRPL50-VC155* constructs. VC was fused to MRPL2 and MRPL50 by linker peptides: GSKQKVMNH. MRPS6 and MRPS16 were fused to VN by linker peptides: GSRSIAT. For the modulation of mtIF3 expression, *AAV-shRNA-ctrl* (Addgene, #85741), and *pcDNA6/V5-HisA* plasmid was modified: *pAAV2-Control-shRNA-TagRFP657*, *pAAV2-mtIF3-shRNA-TagRFP657*, *pcDNA6-5’UTR of mtIF3-3xFLAG-P2A-TagRFP657-3’UTR of mtIF3*, and *pcDNA6-5’UTR of mtIF3-CDS of mtIF3-3xFLAG-P2A-TagRFP657-3’UTR of mtIF3*.

#### Confocal microscopy and image analysis

All the images were taken using a confocal microscope (Zeiss LSM 780). Live cell imaging was performed in a live cell chamber that was maintained at 37°C and 5% CO_2_ by heating instrument. Definite Focus z-correction hardware was used to maintain the z-axis during the time lapse image. Orthogonal projection and image crop were processed in ZEN 3.1 (blue edition). Fluorescence signal intensity was quantified by ImageJ (NIH).

#### Local protein synthesis assay

For local mRNA translation assay, Dendra2 fluorescence protein was conjugated with UTRs of mtIF3 or GAPDH: 5’UTR of mtIF3-Palmitoylation sequence-Dendra2-3’UTR of mtIF3, 5’UTR of mtIF3-CDS of mtIF3-Palmitoylation sequence-Dendra2-3’UTR of mtIF3, and 5’UTR of GAPDH-Palmitoylation sequence-Dendra2-3’UTR of GAPDH. Primary hippocampal neurons were transfected with these vectors at DIV3 by using Lipofectamine 2000 (Invitrogen). 24 hours after transfection, protein synthesis assay was performed. Existing fluorescence of dendra2 (green) was photoconverted into red fluorescence with 405 nm laser for 10 seconds and newly synthesized green signals were measured for 90 minutes with 5 minutes time lapse image. Protein synthesis inhibitor, anisomycin (20 μM, Sigma) was used to confirm that the increased green signal was from *de novo* protein synthesis.

#### Live cell imaging

Before mito-riboBiFC imaging in Neuro2A cells, culture medium was replaced and cells were incubated for 30 minutes. 20 μM of puromycin (Sigma) was applied for 10 minutes. After puromycin treatment, 50 μg/ml of chloramphenicol (Sigma) was sequentially treated. To label mitochondria, mito-mTFP1 was also transfected. Images were acquired using 458, 514nm lasers.

For ATP imaging, primary hippocampal neurons were cultured into microfluidic devices, which were attached on glass bottom dishes. Neurons were transfected with shRNAs at DIV1 and then transfected with overexpression vectors at DIV2. At DIV3, images were acquired using 458, 514, and 633 nm lasers. To fix mitochondrial ribosomes, chloramphenicol was treated for 90 minutes. BDNF or cycloheximide were also treated simultaneously to induce or block local translation and images were taken after 90 minutes. Fluorescent intensity was measured from five mitochondria at the end of axons. The ratio of before and after drug treatment was averaged to measure the degree of mitochondrial translation. For mitochondrial ATP imaging, primary hippocampal neurons were transfected with genetically encoded FRET-based ATP indicator for mitochondria, mito-ATeam1.03, at DIV1 for mtIF3 knockdown study or at DIV2 for mtIF3 overexpression study by using Lipofectamine 2000 (Invitrogen). Images were taken at emission of 475 nm and 527 nm with a 405 nm excitation laser. BDNF or cycloheximide were applied to axonal chamber for 90 minutes. The increased ratio of FRET to CFP before and after drug treatment was calculated.

For mtIF3 localization determination imaging, annotated vector was transfected using PEI in Neuro2A. 250nM of MitoTracker™ Deep Red FM (Invitrogen, M22426) was added to cells and images were acquired after 10 minutes. Images were acquired using 488, 633nm lasers.

#### Western blotting

Cells were lysed by using RIPA buffer (150 mM sodium chloride, 1% Triton X-100, 0.5% sodium deoxycholate and 0.1% sodium dodecyl sulfate). Proteins were separated by SDS-PAGE and transferred to PVDF membranes (Millipore). Membranes were blocked with 5% skim milk in TBST (10 mM Tris, 150 mM NaCl, 0.5% Tween 20) for 30 minutes. For immunoblotting, antibodies against mtIF3 (Sigma, HPA039791, polyclonal) and β-tubulin (Abcam, ab6046, polyclonal) were incubated at 4°C for overnight. Membranes were washed three times for 10 minutes with TBST and horseradish peroxidase-conjugated anti-rabbit secondary antibody (Jackson immunoresearch) was incubated for 1 hour. Membranes were washed three times for 10 minutes with TBST and developed with ECL solution (Bio-Rad).

#### RT-PCR

Total RNA of cell body and axon fractionations was isolated with PicoPure RNA isolation kit (Applied Biosystems). 200 ng of RNA was subjected to RT-PCR by using High Capacity RNA-to-cDNA kit (Life Technologies). The primers used for PCR:

forward 5’-GAGAGCAGATCCACCAGGAG-3’ and
reverse 5’-CTGTTTCCGTCGTCGTCTTT-3’ for mtIF3;
forward 5’-ACCAACTGGGACGACATGGAGAAGA-3’ and
reverse 5’-CGTTGCCAATAGTGATGACCTGGCC-3’ for β-actin;
forward 5’-GGACGACATGGAGAAGATCTGGCAC-3’ and
reverse 5’-CCGGACACCGGAACCGCTCATTG-3’ for γ-actin.

#### Immunostaining

For primary hippocampal neurons and NIH/3T3 cells, cells were rinsed with PBS and fixed with 4% PFA for 10 minutes. Cell were permeabilized with PBST (PBS with 0.2% Triton X-100) for 10 minutes and blocked with 1% BSA in PBST for 30 minutes. Primary antibodies were incubated at 4°C for overnight. Antibodies against Tau1 (Millipore, MAB3420, monoclonal), FLAG (Sigma, F7425, polyclonal), and MTCO1 (Abcam, ab203912, monoclonal) were used for immunostaining. The cells were washed three times with PBS and incubated with Alexa Fluor secondary antibodies (Invitrogen). Then these cells were washed three times with PBS and coverslips were mounted on slide glasses. Images were taken by using LSM780 confocal microscopy. For Neuro2A, cells were washed with ice-cold PBS followed by fixation using 4% PFA/sucrose for 15 minutes. After 3 times washing with PBS, cells were permeabilized with PBST (PBS with 0.5% Triton X-100) for 15 minutes and blocked with 1% BSA in PBS for 1 hour. Primary antibodies were incubated at 4°C for overnight. Cells were washed 3 times with PBS and incubated with Alexa Fluor secondary antibodies (Invitrogen) for 1 hour at room temperature. Cells were washed three times with PBS and coverslips were mounted on slide glasses. Antibodies against GFP (Abcam, ab 6556, polyclonal) and MT-CO1 (Abcam, ab14705, monoclonal) were used for immunostaining.

#### Statistical analysis

Statistical analyses were performed by using Prism software (GraphPad Software) or R (version 3.6.1) with ARTool library (Kay & Wobbrock, 2016; Wobbrock, Findlater, Gergle, & Higgins, 2011). All the values were presented with mean ± SEM. Shapiro-Wilk test for normality (*P*<0.05) or Brown-Forsythe test for equal variance (*P*<0.05) were used to determine the statistical analysis for each dataset. Ordinary two-way ANOVA with Dunnett’s test (repeated measure) or Tukey’s test (non-repeated measure), aligned ranks transformation ANOVA with Wilcoxon signed-rank test (repeated measure) or Wilcoxon rank-sum (non-repeated measure) were used to determine statistical differences between the groups. P < 0.05 was considered as statistically significant. \**P* < 0.05, \*\**P* < 0.01, \*\*\**P* < 0.001, \*\*\*\**P* < 0.0001.

## Acknowledgements

We wish to dedicate this work to the life and career of our beloved mentor and colleague, Dr. Kyung-Tai Min, who passed away recently. This work was supported by Basic Science Research Program through the National Research Foundation of Korea (NRF) funded by the Ministry of Science and ICT (2016R1A3B1905982 to K.M., 2020R1A2C1005492 to J.-I.K.). This work was also supported by a grant from the Suh Kyungbae Foundation (SUHF-17020101 to C.L.).

## Author contribution

Conceptualization, D.P. and K.M.; Methodology, S.L., D.P., C.L., J.-I.K. and K.M.; Investigation, S.L. and D.P.; Interpretation of Data, S.L., D.P., C.L., J.-I.K. and K.M.; Writing – Original Draft, S.L., D.P. and K.M.; Writing – Review & Editing, S.L., D.P., C.L. and J.-I.K.; Funding Acquisition, C.L., J.-I.K. and K.M.; Supervision, C.L., J.-I.K. and K.M.

## Supplementary figure legend

**Figure 1-figure supplement 1.**
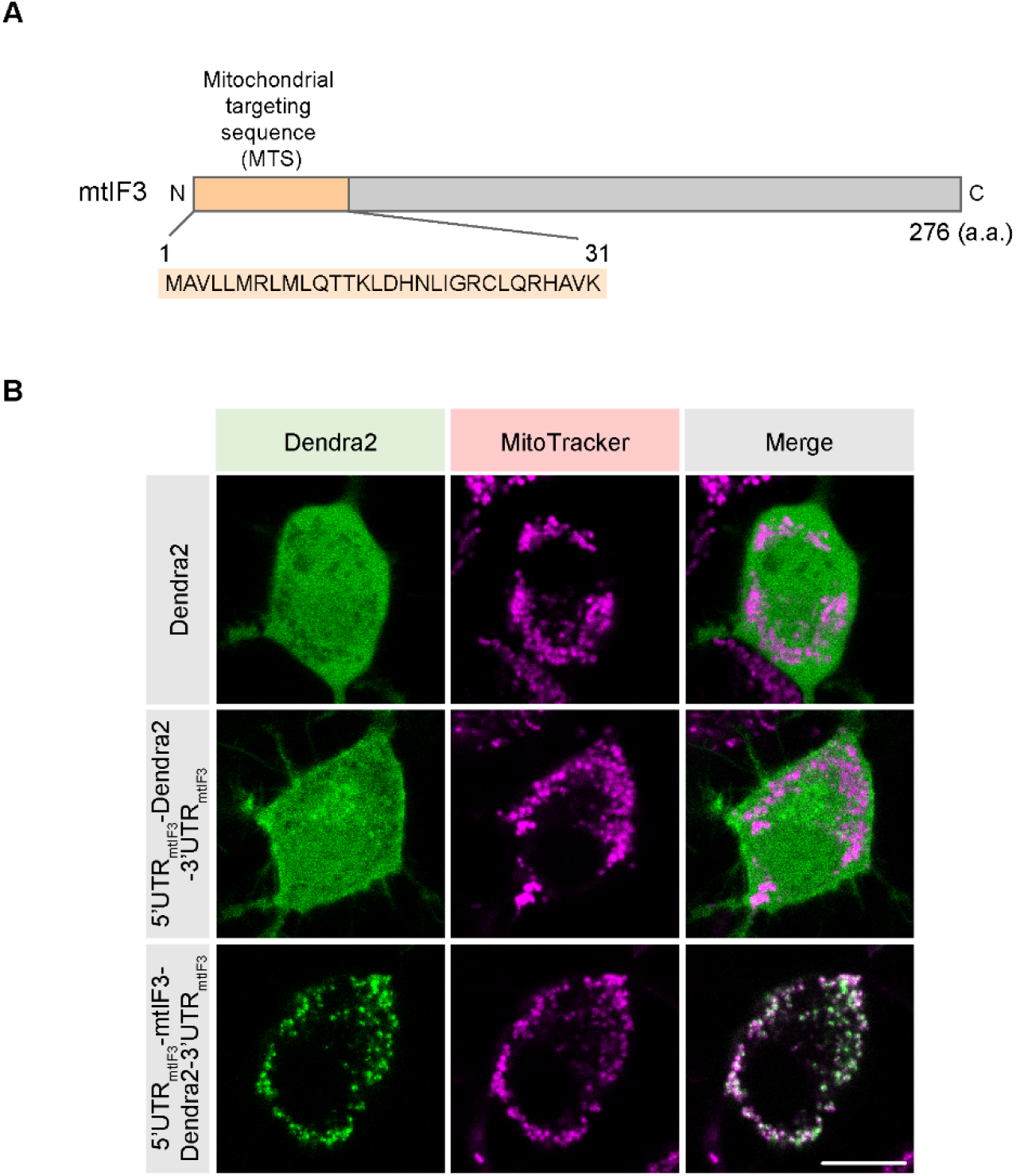
mtIF3-Dendra2 is localized to mitochondria. **(A)** Schematic illustration of mtIF3 coding sequence. mtIF3 has mitochondrial targeting sequence in N-terminal domain (131 a.a.) (Koc & Spremulli, 2002). **(B)** Neuro2A cells were transfected with Dendra2 vectors. Mitochondria were labeled with MitoTracker deep red dye. CDS of mtIF3 led to mitochondrial localization of Dendra2 (scale bar, 10 μm).

**Figure 2-figure supplement 1.**
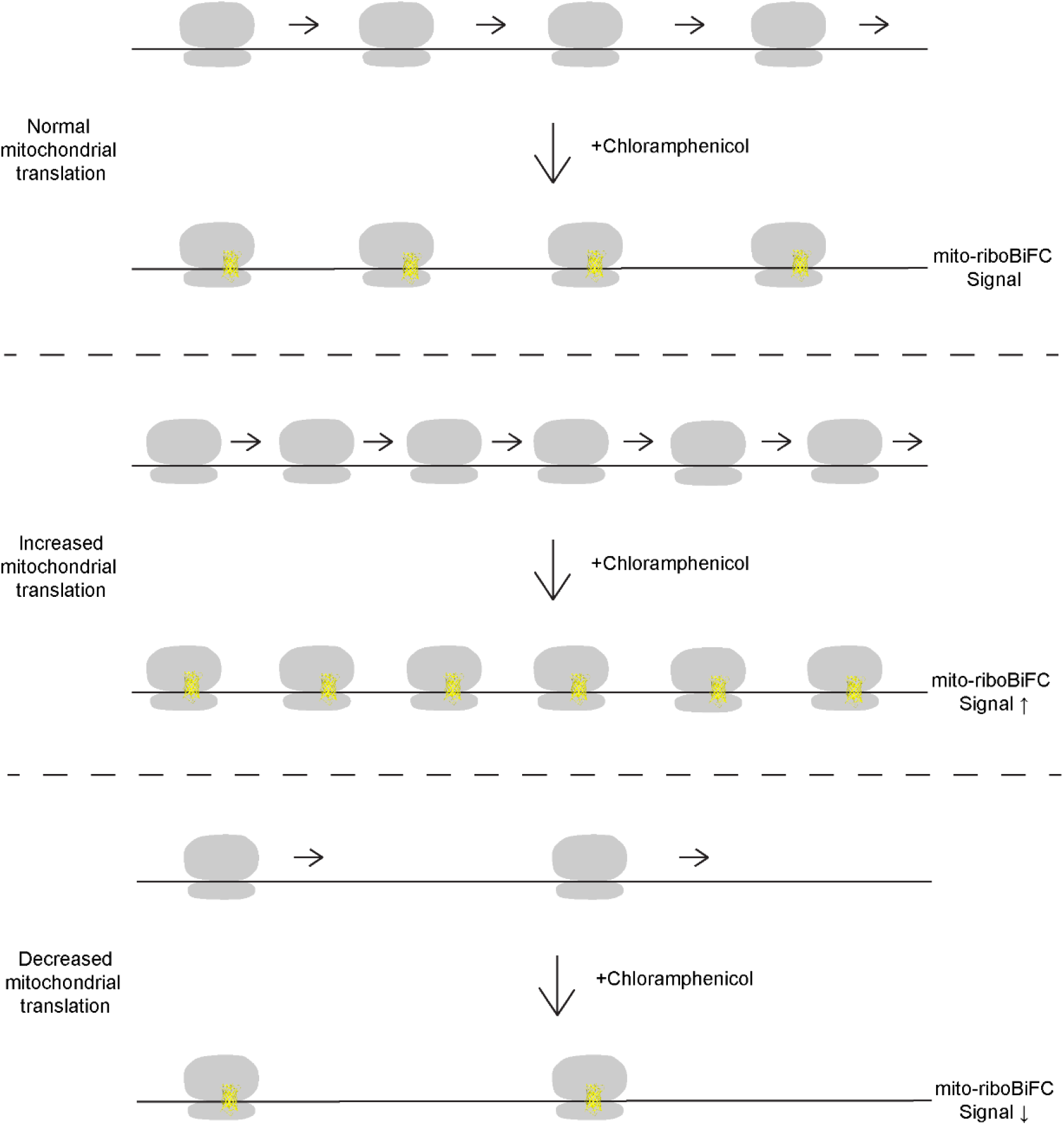
Schematic illustration of mito-riboBiFC analysis. Translating ribosomal complex exhibits active dynamics. During the elongation, ribosomal subunits consistently rotate, which results in low intensity mito-riboBiFC. To freeze translating mitochondrial ribosomes, we treated chloramphenicol that inhibits the formation of peptide bond. Non-rotated ribosomal complex is expected to show high BiFC signal. 90 minutes after the treatment of chloramphenicol, we compared the intensity of mito-riboBiFC before and after chloramphenicol treatment. Because highly translating mRNA binds to more ribosomes, we could detect higher signal increase in actively translating mRNA.

**Figure 3-figure supplement 1.**
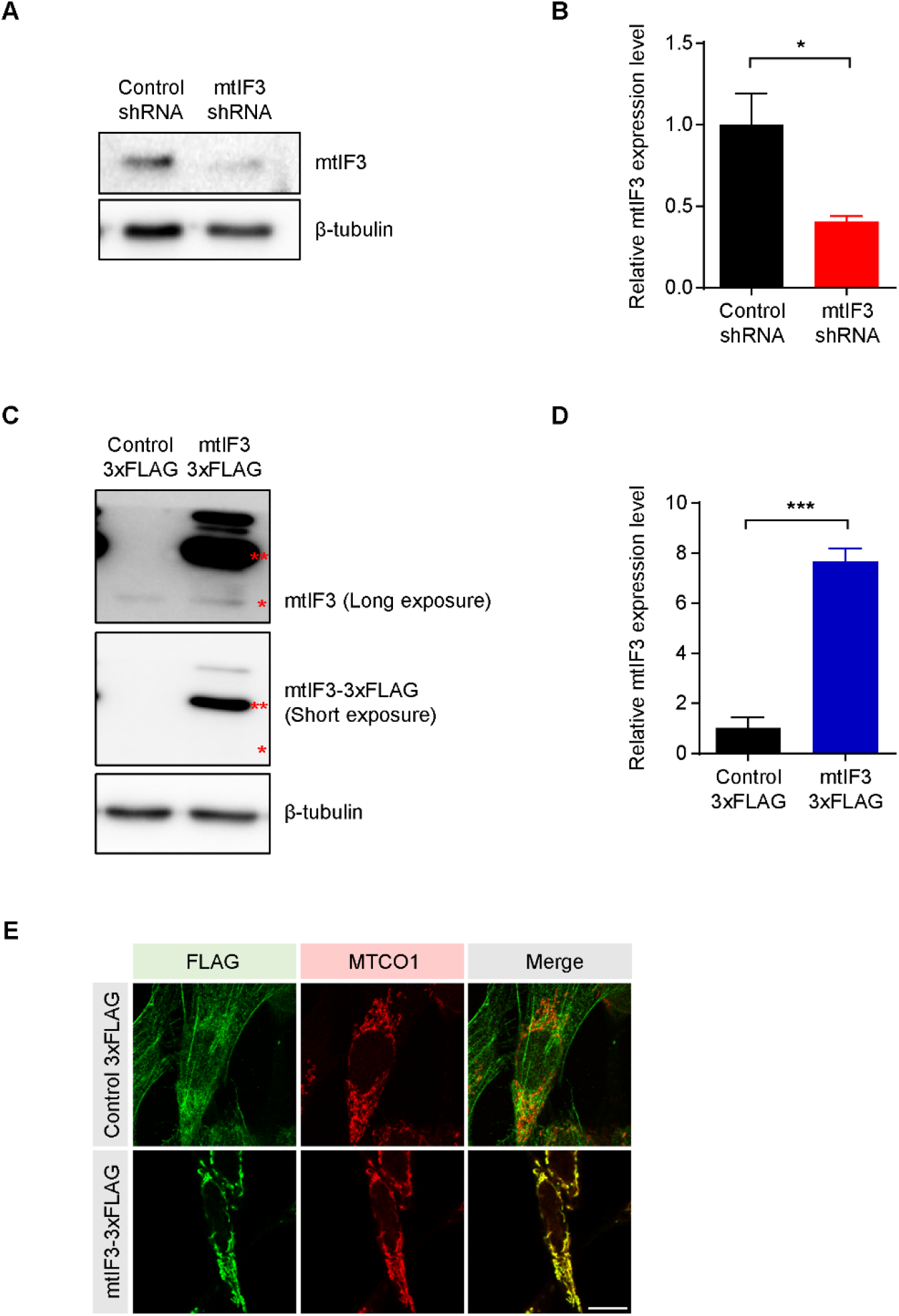
Modulation of mtIF3 expression level. **(A, B)** mtIF3 level was reduced by RNA interference with shRNA. By performing western blot, we verified the successful reduction of mtiF3 48 hours after incubation of shRNA in NIH/3T3 cells. The level of mtIF3 was decreased about 50% compared with control group. The mtIF3 expression level was normalized to β-tubulin level. N = 5 independent experiments. (**C, D**) mtIF3 was upregulated by overexpression vector in NIH/3T3 cells. Overexpression vectors contain tandems of FLAG epitope tag. Western blot results showed the expression of mtIF3 overexpression vector. Single red asterisk indicates endogenous mtIF3 and double red asterisk indicates exogenous mtIF3 that is conjugated with FLAG tag. The level of overexpressed mtIF3 was about 8 times higher than endogenous mtIF3 level. N = 3 independent experiments. (**E**) Immunofluorescence images demonstrating the localization of overexpressed mtIF3 to mitochondria in NIH/3T3 cells. FLAG staining detected the transfected overexpression or control vectors. Mitochondria were marked by MTCO1 staining (scale bar, 10 μm). The values were presented as mean ± SEM and statistical significance was analyzed by unpaired t-test. \**P* < 0.05, \*\*\**P* < 0.001.

**Figure 4-figure supplement 1.**
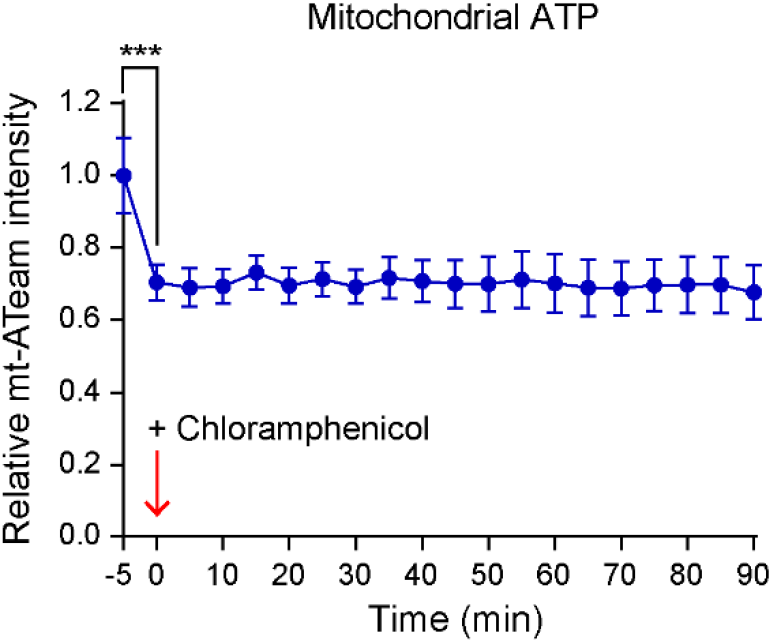
Blocking mitochondrial translation decreases mitochondrial ATP level. Treatment of chloramphenicol caused the rapid decrease of FRET intensity. Primary hippocampal neurons were transfected with Mt-ATeam1.03 at DIV2 and images were taken every 5 minutes for 90 minutes at DIV3. The values are presented as mean ± SEM and statistical significance was tested between 5 minutes before treatment and 0 minute after treatment using paired t-test. N = 10 cells from 3 independent experiments. \*\*\**P* < 0.001.

**Figure 5-figure supplement 1.**
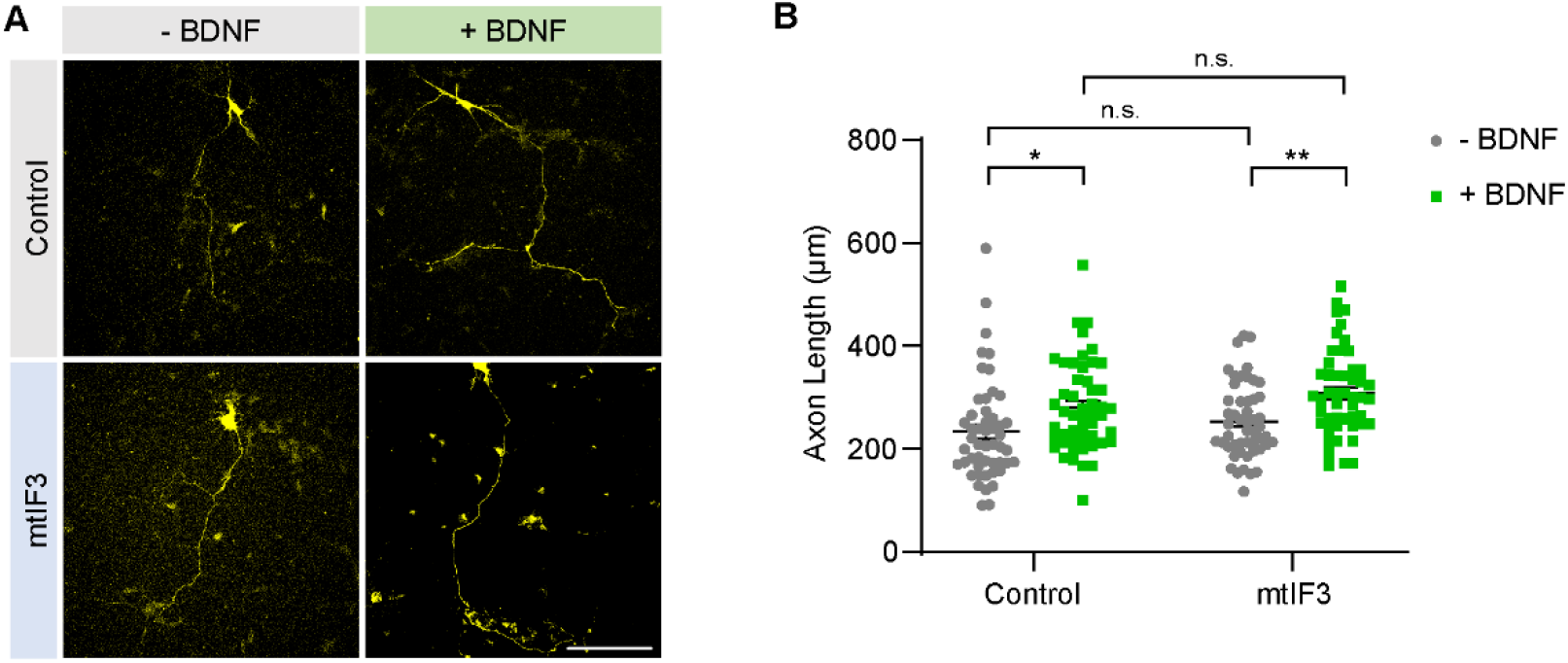
mtIF3 overexpression is not enough to promote axon extension, but BDNF treatment still facilitates axon growth. **(A)** Representative images of primary neurons expressing mtIF3 overexpression vectors. Hippocampal neurons were transfected with control and overexpression vectors at DIV2 and BDNF was also treated simultaneously. Images were taken at DIV3. Transfection of overexpression vector was confirmed by TagRFP657 expression. Hippocampal neurons were identified by their morphology (scale bar, 100 μm). **(B)** Quantification of axonal length. Overexpression of mtIF3 had no additional effect on axon growth, while BDNF treatment still promoted axonal length. Data represent mean ± SEM (N = 50 neurons from 5 independent experiments). Aligned ranks transformation ANOVA detected no significant interaction effects of mtIF3 overexpression and drug treatment. n.s., not significant; \**P* < 0.05, \*\**P* < 0.01, as determined by Wilcoxon rank-sum test.

## References

Al-Jubran, K., Wen, J., Abdullahi, A., Roy Chaudhury, S., Li, M., Ramanathan, P., … Brogna, S. (2013). Visualization of the joining of ribosomal subunits reveals the presence of 80S ribosomes in the nucleus. RNA, 19(12), 1669–1683. doi:10.1261/rna.038356.113

Amunts, A., Brown, A., Toots, J., Scheres, S. H. W., & Ramakrishnan, V. (2015). The structure of the human mitochondrial ribosome. Science, 348(6230), 95–98. doi:10.1126/science.aaa1193

Aschrafi, A., Kar, A. N., Gale, J. R., Elkahloun, A. G., Vargas, J. N., Sales, N., … Kaplan, B. B. (2016). A heterogeneous population of nuclear-encoded mitochondrial mRNAs is present in the axons of primary sympathetic neurons. Mitochondrion, 30, 18–23. doi:10.1016/j.mito.2016.06.002

Aschrafi, A., Natera-Naranjo, O., Gioio, A. E., & Kaplan, B. B. (2010). Regulation of axonal trafficking of cytochrome c oxidase IV mRNA. Molecular and Cellular Neuroscience, 43(4), 422–430.

Barsh, G. S., Lagouge, M., Mourier, A., Lee, H. J., Spåhr, H., Wai, T., … Larsson, N.-G. (2015). SLIRP Regulates the Rate of Mitochondrial Protein Synthesis and Protects LRPPRC from Degradation. PLOS Genetics, 11(8), e1005423. doi:10.1371/journal.pgen.1005423

Chatenay-Lapointe, M., & Shadel, G. S. (2011). Repression of mitochondrial translation, respiration and a metabolic cycle-regulated gene, SLF1, by the yeast Pumilio-family protein Puf3p. PLoS One, 6(5), e20441. doi:10.1371/journal.pone.0020441

Christian, B. E., & Spremulli, L. L. (2009). Evidence for an Active Role of IF3mt in the Initiation of Translation in Mammalian Mitochondria. Biochemistry, 48(15), 3269–3278. doi:10.1021/bi8023493

Couvillion, M. T., Soto, I. C., Shipkovenska, G., & Churchman, L. S. (2016). Synchronized mitochondrial and cytosolic translation programs. Nature, 533(7604), 499–503. doi:10.1038/nature18015

Estell, C., Stamatidou, E., El-Messeiry, S., & Hamilton, A. (2017). In situ imaging of mitochondrial translation shows weak correlation with nucleoid DNA intensity and no suppression during mitosis. J Cell Sci, 130(24), 4193–4199. doi:10.1242/jcs.206714

Gale, J. R., Aschrafi, A., Gioio, A. E., & Kaplan, B. B. (2018). Nuclear-Encoded Mitochondrial mRNAs: A Powerful Force in Axonal Growth and Development. Neuroscientist, 24(2), 142–155. doi:10.1177/1073858417714225

Greber, B. J., Bieri, P., Leibundgut, M., Leitner, A., Aebersold, R., Boehringer, D., & Ban, N. (2015). The complete structure of the 55S mammalian mitochondrial ribosome. Science, 348(6232), 303–308. doi:10.1126/science.aaa3872

Gumy, L. F., Yeo, G. S., Tung, Y.-C. L., Zivraj, K. H., Willis, D., Coppola, G., … Fawcett, J. W. (2011). Transcriptome analysis of embryonic and adult sensory axons reveals changes in mRNA repertoire localization. Rna, 17(1), 85–98.

Hafner, A. S., Donlin-Asp, P. G., Leitch, B., Herzog, E., & Schuman, E. M. (2019). Local protein synthesis is a ubiquitous feature of neuronal pre- and postsynaptic compartments. Science, 364(6441). doi:10.1126/science.aau3644

Han, S. M., Baig, H. S., & Hammarlund, M. (2016). Mitochondria Localize to Injured Axons to Support Regeneration. Neuron, 92(6), 1308–1323. doi:10.1016/j.neuron.2016.11.025

Hillefors, M., Gioio, A. E., Mameza, M. G., & Kaplan, B. B. (2007). Axon viability and mitochondrial function are dependent on local protein synthesis in sympathetic neurons. Cell Mol Neurobiol, 27(6), 701–716. doi:10.1007/s10571-007-9148-y

Hu, C.-D., Chinenov, Y., & Kerppola, T. K. (2002). Visualization of interactions among bZIP and Rel family proteins in living cells using bimolecular fluorescence complementation. Molecular cell, 9(4), 789–798.

Imamura, H., Nhat, K. P., Togawa, H., Saito, K., Iino, R., Kato-Yamada, Y., … Noji, H. (2009). Visualization of ATP levels inside single living cells with fluorescence resonance energy transfer-based genetically encoded indicators. Proc Natl Acad Sci U S A, 106(37), 15651–15656. doi:10.1073/pnas.0904764106

Ingolia, N. T., Lareau, L. F., & Weissman, J. S. (2011). Ribosome profiling of mouse embryonic stem cells reveals the complexity and dynamics of mammalian proteomes. Cell, 147(4), 789–802. doi:10.1016/j.cell.2011.10.002

Jung, H., Yoon, B. C., & Holt, C. E. (2012). Axonal mRNA localization and local protein synthesis in nervous system assembly, maintenance and repair. Nat Rev Neurosci, 13(5), 308–324. doi:10.1038/nrn3210

Kaplan, B. B., Gioio, A. E., Hillefors, M., & Aschrafi, A. (2009). Axonal protein synthesis and the regulation of local mitochondrial function. Results Probl Cell Differ, 48, 225–242. doi:10.1007/400_2009_1

Kay, M., & Wobbrock, J. (2016). ARTool: aligned rank transform for nonparametric factorial ANOVAs. R package version 0.10, 2.

Koc, E. C., & Spremulli, L. L. (2002). Identification of mammalian mitochondrial translational initiation factor 3 and examination of its role in initiation complex formation with natural mRNAs. J Biol Chem, 277(38), 35541–35549. doi:10.1074/jbc.M202498200

Kuzniewska, B., Cysewski, D., Wasilewski, M., Sakowska, P., Milek, J., Kulinski, T. M., … Dziembowska, M. (2020). Mitochondrial protein biogenesis in the synapse is supported by local translation. EMBO Rep, e48882. doi:10.15252/embr.201948882

Lee, S., Wang, W., Hwang, J., Namgung, U., & Min, K. T. (2019). Increased ER-mitochondria tethering promotes axon regeneration. Proc Natl Acad Sci U S A, 176(32), 16074–16079. doi:10.1073/pnas.1818830116

Natera-Naranjo, O., Kar, A. N., Aschrafi, A., Gervasi, N. M., Macgibeny, M. A., Gioio, A. E., & Kaplan, B. B. (2012). Local translation of ATP synthase subunit 9 mRNA alters ATP levels and the production of ROS in the axon. Molecular and Cellular Neuroscience, 49(3), 263–270.

Niescier, R. F., Kwak, S. K., Joo, S. H., Chang, K. T., & Min, K. T. (2016). Dynamics of Mitochondrial Transport in Axons. Front Cell Neurosci, 10, 123. doi:10.3389/fncel.2016.00123

Park, D., Lee, S., & Min, K.-T. (2020). Techniques for investigating mitochondrial gene expression. BMB Reports, 53(1), 3–9. doi:10.5483/BMBRep.2020.53.1.272

Rangaraju, V., Lauterbach, M., & Schuman, E. M. (2019). Spatially Stable Mitochondrial Compartments Fuel Local Translation during Plasticity. Cell, 176(1-2), 73–84 e15. doi:10.1016/j.cell.2018.12.013

Richter-Dennerlein, R., Oeljeklaus, S., Lorenzi, I., Ronsor, C., Bareth, B., Schendzielorz, A. B., … Dennerlein, S. (2016). Mitochondrial Protein Synthesis Adapts to Influx of Nuclear-Encoded Protein. Cell, 167(2), 471–483 e410. doi:10.1016/j.cell.2016.09.003

Richter, U., Lahtinen, T., Marttinen, P., Suomi, F., & Battersby, B. J. (2015). Quality control of mitochondrial protein synthesis is required for membrane integrity and cell fitness. Journal of Cell Biology, 211(2), 373–389. doi:10.1083/jcb.201504062

Robida, A. M., & Kerppola, T. K. (2009). Bimolecular fluorescence complementation analysis of inducible protein interactions: effects of factors affecting protein folding on fluorescent protein fragment association. J Mol Biol, 394(3), 391–409. doi:10.1016/j.jmb.2009.08.069

Rose, R. H., Briddon, S. J., & Holliday, N. D. (2010). Bimolecular fluorescence complementation: lighting up seven transmembrane domain receptor signalling networks. Br J Pharmacol, 159(4), 738–750. doi:10.1111/j.1476-5381.2009.00480.x

Rudler, D. L., Hughes, L. A., Perks, K. L., Richman, T. R., Kuznetsova, I., Ermer, J. A., … Hool, L. C. (2019). Fidelity of translation initiation is required for coordinated respiratory complex assembly. Science advances, 5(12), eaay2118.

Saxton, W. M., & Hollenbeck, P. J. (2012). The axonal transport of mitochondria. J Cell Sci, 125(Pt 9), 2095–2104. doi:10.1242/jcs.053850

Sheng, Z. H. (2017). The Interplay of Axonal Energy Homeostasis and Mitochondrial Trafficking and Anchoring. Trends Cell Biol, 27(6), 403–416. doi:10.1016/j.tcb.2017.01.005

Shigeoka, T., Jung, H., Jung, J., Turner-Bridger, B., Ohk, J., Lin, J. Q., … Holt, C. E. (2016). Dynamic Axonal Translation in Developing and Mature Visual Circuits. Cell, 166(1), 181–192. doi:10.1016/j.cell.2016.05.029

Shigeoka, T., Koppers, M., Wong, H. H., Lin, J. Q., Cagnetta, R., Dwivedy, A., … Holt, C. E. (2019). OnSite Ribosome Remodeling by Locally Synthesized Ribosomal Proteins in Axons. Cell Rep, 29(11), 3605–3619 e3610. doi:10.1016/j.celrep.2019.11.025

Smits, P., Smeitink, J., & van den Heuvel, L. (2010). Mitochondrial Translation and Beyond: Processes Implicated in Combined Oxidative Phosphorylation Deficiencies. Journal of Biomedicine and Biotechnology, 2010, 737385. doi:10.1155/2010/737385

Spillane, M., Ketschek, A., Merianda, T. T., Twiss, J. L., & Gallo, G. (2013). Mitochondria coordinate sites of axon branching through localized intra-axonal protein synthesis. Cell Rep, 5(6), 1564–1575. doi:10.1016/j.celrep.2013.11.022

Taylor, A. M., Blurton-Jones, M., Rhee, S. W., Cribbs, D. H., Cotman, C. W., & Jeon, N. L. (2005). A microfluidic culture platform for CNS axonal injury, regeneration and transport. Nat Methods, 2(8), 599–605. doi:10.1038/nmeth777

Vaarmann, A., Mandel, M., Zeb, A., Wareski, P., Liiv, J., Kuum, M., … Kaasik, A. (2016). Mitochondrial biogenesis is required for axonal growth. Development, 143(11), 1981–1992. doi:10.1242/dev.128926

Wang, W., Rai, A., Hur, E. M., Smilansky, Z., Chang, K. T., & Min, K. T. (2016). DSCR1 is required for both axonal growth cone extension and steering. J Cell Biol, 213(4), 451–462. doi:10.1083/jcb.201510107

Wobbrock, J. O., Findlater, L., Gergle, D., & Higgins, J. J. (2011). The aligned rank transform for nonparametric factorial analyses using only anova procedures. Paper presented at the Proceedings of the SIGCHI conference on human factors in computing systems.

Yoon, B. C., Jung, H., Dwivedy, A., O’Hare, C. M., Zivraj, K. H., & Holt, C. E. (2012). Local translation of extranuclear lamin B promotes axon maintenance. Cell, 148(4), 752–764. doi:10.1016/j.cell.2011.11.064

Zhou, B., Yu, P., Lin, M. Y., Sun, T., Chen, Y., & Sheng, Z. H. (2016). Facilitation of axon regeneration by enhancing mitochondrial transport and rescuing energy deficits. J Cell Biol, 214(1), 103–119. doi:10.1083/jcb.201605101

